# The His-tag as a decoy modulating preferred orientation in cryoEM

**DOI:** 10.1101/2020.09.22.309005

**Authors:** Raquel Bromberg, Yirui Guo, Daniel Plymire, Tabitha Emde, Maciej Puzio, Dominika Borek, Zbyszek Otwinowski

**Affiliations:** Department of Biophysics, The University of Texas Southwestern Medical Center, 5323 Harry Hines Blvd., Dallas, TX, 75390, United States; Ligo Analytics, 2207 Chunk Ct, Dallas, TX, 75206-0057, United States; Department of Biochemistry, The University of Texas Southwestern Medical Center, 5323 Harry Hines Blvd., Dallas, TX, 75390, United States

**Author notes:** Ligo Analytics, 2207 Chunk Ct, Dallas, TX 75206-0057, United States. **Funding information** The National Institute of Allergy and Infectious Diseases (NIAID), National Institutes of Health, Department of Health and Human Services (contract No. HHSN272201700060C to Zbyszek Otwinowski); The National Institute of General Medical Sciences (NIGMS), National Institutes of Health, (grant No. R21GM126406 to Dominika Borek; grant No. R01GM117080 to Zbyszek Otwinowski; grant No. R01GM118619 to Zbyszek Otwinowski; grant No. R43GM137671 to Yirui Guo); Department of Energy (grant No. DESC0019600 to Yirui Guo); The National Institute of Allergy and Infectious Diseases (NIAID), National Institutes of Health (grant No. P01AI120943 to Dominika Borek, Zbyszek Otwinowski).

**Keywords:** cryo-electron microscopy single particle reconstruction (cryoEM SPR), water-air interface, His-tag, preferred orientation

## Abstract

The His-tag is a widely used affinity tag that facilitates purification by means of affinity chromatography of recombinant proteins for functional and structural studies. We show here that His-tag presence affects how coproheme decarboxylase interacts with the water-air interface during grid preparation for cryoEM. Depending on His-tag presence or absence, we observe significant changes in patterns of preferred orientation. The analysis of particle orientations suggests that His-tag presence can mask the hydrophobic patches on a protein’s surface that mediate the interactions with the water-air interface, while the hydrophobic linker between a His-tag and the coding sequence of the protein may enhance other interactions with water-air interface. Our observations suggest that tagging, including rational design of the linkers between an affinity tag and a protein of interest, offer a promising approach to modulating interactions with the water-air interface.

**Synopsis:** A His-tag affects the interactions of particles with the water-air interface in cryo-electron microscopy (cryoEM) single particle reconstruction (SPR), and thus may be used to modulate these interactions, including inducing changes in patterns of preferred orientation.

## 1. Introduction

cryoEM single particle reconstruction (SPR) datasets consist of images of macromolecular particles dispersed in a thin layer of amorphous ice formed over a thin supporting material that is attached to a metal grid. In recent years, there have been several studies on the angular and positional orientations of particles within the ice layer in cryoEM SPR and on the interactions of particles with the water-air interface during cryo-cooling. In these studies, the kinetic parameters, variations in mechanical support, and chemical modifications to the support were explored. A number of strategies helping to keep biological samples in their optimal state have been proposed (D’Imprima *et al*., 2019, Glaeser, 2018, Noble, Wei, *et al*., 2018, Drulyte *et al*., 2018).

The water-air interface has been shown (Noble, Dandey, *et al*., 2018, D’Imprima *et al*., 2019, Chen *et al*., 2019) to have damaging influence due to the extremely high hydrophobicity of air (van Oss *et al*., 2005). Many macromolecular particles have on their surface hydrophobic patches which are strongly attracted to the water-air interface (Glaeser & Han, 2017, Taylor & Glaeser, 2008), and diffusion allows particles to reach the interface within milliseconds, with partial or full unfolding frequently following initial binding (D’Imprima *et al*., 2019, Noble, Dandey, *et al*., 2018, Glaeser & Han, 2017). Even if molecules do not unfold, they may become preferentially oriented (Noble, Dandey, *et al*., 2018, Han *et al*., 2012, Tan *et al*., 2017, Chen *et al*., 2019). The binding to the water-air interface has also a significant advantage in reducing particle exclusion from thin ice, which is highly preferred for obtaining high quality data. Therefore, interactions between particles and the water-air interface have a complex impact upon experimental results.

Preferred orientation hinders cryoEM SPR reconstruction in two ways (Radermacher, 1988). Firstly, the preferred orientation results in a systematic lack of information regarding some orientations (the so-called missing cone effect), which leads to uneven coverage of reciprocal space and thus can be considered analogous to anisotropic diffraction in X-ray crystallography. In addition, the lack of a large group of orientations affects the convergence of the computational procedures used in 3D reconstruction for cryoEM SPR. Particularly for macromolecular systems having molecular mass lower than ~150 kDa, systematic misalignment of particles will result in introducing bias and artifacts to the reconstruction (Barth *et al*., 1989, Naydenova & Russo, 2017, Tan *et al*., 2017, Noble, Dandey, *et al*., 2018, Radermacher, 1988).

Several approaches have been proposed to alleviate the problem of damage induced by the water-air interface or of the preferred orientation induced by it. They include using surfactants (Chen *et al*., 2019, Glaeser *et al*., 2016, Frederik *et al*., 1989) to saturate the surface at the water-air interface or the surface of the support (Russo & Passmore, 2014), a graphene oxide or graphene supports to prevent interactions the water-air interface (Pantelic *et al*., 2010, Pantelic *et al*., 2011, Wang, Liu, *et al*., 2020, Wang, Yu, *et al*., 2020). These supports may also be chemically modified to promote specific, high-affinity interactions with particles (Llaguno *et al*., 2014, Kelly *et al*., 2010a, Kelly *et al*., 2008, Kelly *et al*., 2010b, Han *et al*., 2012, Benjamin, Wright, Hyun, *et al*., 2016, Benjamin, Wright, Bolton, *et al*., 2016, Crucifix *et al*., 2004, Wang, Liu, *et al*., 2020, Wang, Yu, *et al*., 2020, Yeates *et al*., 2020). Finally, new, fast approaches for depositing samples on a grid have been developed (Arnold *et al*., 2017, Schmidli *et al*., 2018, Dandey *et al*., 2020, Dandey *et al*., 2018, Wei *et al*., 2018, Ravelli *et al*., 2019). One of the strategies for changing interactions of macromolecules with a support, and with the water-air interface, is to chemically modify the molecule itself without changing the chemical properties of the support. Affinity tags chemically modify a protein and have been used in sample preparation for cryoEM in the context of affinity purification (Wang, Liu, *et al*., 2020, Wang, Yu, *et al*., 2020, Benjamin, Wright, Bolton, *et al*., 2016, Benjamin, Wright, Hyun, *et al*., 2016), but as we observed, tag presence can modify interactions with the water-air interface even without the use of affinity grids.

The His-tag is an affinity tag used widely to facilitate purification of recombinant proteins to homogeneity (Hochuli *et al*., 1988). Such proteins, with 6 to 10 consecutive histidines inside the tag which is introduced at their N- or C-termini, are overexpressed, with the tag facilitating purification by immobilized metal ion affinity chromatography (IMAC) (Wong *et al*., 1991). After purification, the His-tag is usually cleaved with a protease which recognizes the specific site. Several proteolytic enzymes are used for this purpose, e.g. thrombin, factor Xa, Tobacco Etch Virus (TEV)-protease, carboxypeptidase A. TEV-protease is a common choice because its classical cleavage site ENLYFQ(G/S) is very specific; the cleavage is performed after a glutamine residue, with TEV cleavage generating only a single amino acid addition if the His-tag was used at the N-terminal end of the expressed protein. When used for this purpose, TEV protease is usually expressed with an uncleavable His-tag (Raran-Kurussi *et al*., 2017). After the first round of purification, when a His-tagged protein of interest is separated from untagged proteins using IMAC, His-tag labelled TEV protease is added to the protein of interest. TEV protease cleaves His-tags from the protein of interest, and this cleavage is followed by a second round of IMAC to capture both the His-tag labelled TEV protease and also molecules with uncleaved His-tags, while the His-tag free protein is collected in the flow-through fractions. This procedure is frequently followed by an additional chromatography step, e.g. native gel filtration, to assure that the protein is properly folded and in its proper oligomeric state.

The presence of the His-tag in crystallization is usually considered an obstacle, as it is usually connected to the protein of interest by a flexible linker and the increased flexibility may interfere with crystallization success (Majorek *et al*., 2014). However, in the case of cryoEM SPR, this does not necessarily pose a problem. Here we show and discuss how the His-tag affects interactions between labelled proteins and the water-air interface, which results in the modulation of particle preferred orientations.

## 2. Methods

Coproheme decarboxylases (formerly HemQ) from *Geobacillus stearothermophilus* was one of the targets of the Midwest Center for Structural Genomics (MCSG; target APC35880). We solved its X-ray crystallographic structure in 2004 (PDB code: 1T0T.pdb), while others determined its function later (Dailey & Gerdes, 2015, Milazzo *et al*., 2019, Pfanzagl *et al*., 2018). We recently characterized this protein using cryoEM SPR (EMPIAR-10363, EMD-21373 and EMPIAR-10362, EMD-21376) (Bromberg *et al*., 2020) and noticed during cryoEM experiments that batches of the protein purified at different times showed different patterns of preferred orientation during cryoEM SPR data collection. This observation prompted the analysis presented here.

### 2.1. Protein expression and purification

Coproheme decarboxylase from *Geobacillus stearothermophilus* (TOT) is encoded by the GYMC52_3505 plasmid which is available from the DNASU Plasmid Repository (https://dnasu.org/). The open reading frame of the protein is cloned in the pMCSG7 vector containing Tobacco Etch Virus (TEV) cleavable N-terminal His6-tag (Stols *et al*., 2002).

His-tagged Tobacco Etch Virus (TEV) protease encoded by the pMHTDelta238 plasmid is available from the DNASU Plasmid Repository (https://dnasu.org/). It expresses a mutated and truncated form of the TEV protease as an N-terminal fusion to MBP-His7 (Raran-Kurussi *et al*., 2017), in which the MBP fusion is removed in vivo by autocleavage, leaving His7-TEV.

Our protein expression and purification followed a previously established protocol (Kim *et al*., 2011) with modifications that we describe here in detail.

The GYMC52_3505 and pMHTDelta238 plasmids were transformed to Rosetta2(DE3)pLysS competent cells (Cat. No. 71401-3, EMD Millipore). Transformation reactions were spread on LB plates with ampicillin (Amp) at 200 μg/ml and chloramphenicol (Cam) at 37 μg/ml for GYMC52_3505, and with kanamycin (Kan) at 25 μg/ml and Cam at 37 μg/ml for pMHTDelta238. Single colonies grown on selective LB plates were used to initiate 3 ml or 25 ml liquid media cultures of Luria Broth (LB) with 200 μg/ml Amp and 37 μg/ml Cam for coproheme decarboxylase, and with 50 μg/ml Kan and 37 μg/ml Cam for TEV. All cultures were grown overnight in an incubated shaker (225 rpm, 37 °C). The next morning, these cultures were used to seed larger volume cultures at the ratio 1:1000. 1 L or 6 L LB cultures, with appropriate antibiotics, were grown at 37 °C with 225 rpm shaking until OD600 of ~1.0, when the expression was induced by adding Isopropyl β-d-1-thiogalactopyranoside (IPTG) to the final concentration of 1 mM. The temperature in the shaker was decreased to 28 °C for coproheme decarboxylase and 20 °C for TEV, and cultures were grown overnight at these temperatures with shaking at 225 rpm. The cultures were centrifuged, and pellets were further processed and purified with slightly different protocols.

The bacterial pellets were resuspended in the lysis buffer (50 mM sodium phosphate pH 7.5, 300 mM NaCl buffer) at ~5 ml of the buffer per 1 g of the bacterial pellet. One tablet of cOmplete™ Protease Inhibitor Cocktail (Roche) was added to lysate. The bacterial suspension was then sonicated on ice for 15 minutes [30 × (30/30 s on/off), at 50 % amplitude]. The lysate was centrifuged at 45,000×g for 30 minutes. The supernatant was retained and clarified by filtration through the 0.8 μm filter Millex-AA Syringe Filter Uni (MCE/blue colour). The clarified lysate was applied on the column containing 3-4 ml of Talon resin preequilibrated with a binding buffer consisting of 50 mM sodium phosphate pH 7.5, 300 mM NaCl buffer. After the lysate flew gravitationally through the column, ~100 ml of the binding buffer was applied to remove non-specifically bound proteins. The protein was eluted from the column in 1.5-2 ml fractions by applying ~15 ml of the elution buffer consisting of 50 mM sodium phosphate pH 7.5, 300 mM NaCl buffer, 150 mM imidazole. The aliquot of lysate as well as aliquots of the flow-through fraction, a fraction from the wash with binding buffer and a fraction from the wash with wash buffer, and each of the elution fractions were analysed with the SDS-PAGE. The fractions containing coproheme decarboxylase or TEV proteins were pooled. Purified according to the above protocol, His-tagged TEV protease (~1 mg) was added to pooled fractions of coproheme decarboxylase proteins and dialyzed 24-48 hours against the binding buffer. Then dialysate was applied to 5 ml of Talon resin equilibrated with the binding buffer and flow-through was collected. The flow-through was concentrated to ~2 ml, filtered through a 0.2 μm centrifugal filter and applied to a Superdex^®^ 200 10/300 column run with an AKTA Pure system at a rate of 1 mL/minute in 50 mM HEPES pH 7.5, 100 mM NaCl. Coproheme decarboxylase eluted at ~12 ml, in agreement with the molecular mass of a pentamer (144 kDa). The fractions were collected, analyzed on the SDS-PAGE and concentrated with Amicon Ultra centrifugal concentrator with a 10 kDa molecular mass cutoff (Millipore) to the desired concentrations (20-30 mg/ml), which were assessed by measuring UV absorbance at 280 nm. In addition, centrifugal filtration was accompanied with buffer exchange to 50 mM HEPES pH 7.5 100 mM NaCl. For partially cleaved batches of the protein, we did not use the second IMAC step, and instead applied the sample after TEV digestion directly to the gel filtration column with the assumption that it contained a mixture of tagged states in the pentamer.

### 2.2. Preparation of grids, cryoEM data collection and data analysis

The purified coproheme decarboxylase samples were used to prepare grids. We used gold Quantifoil R 1.2/1.3 grids. The grids were glow discharged 90 s at 30 mA with a PELCO easiGlow™ Glow Discharge Cleaning System to obtain a hydrophilic surface. The glow-discharged grids were used to prepare vitrified samples with the Thermo Scientific Vitrobot Mark IV System. We applied 3 μl of purified protein to the glow-discharged surface of the grid at 4 °C, 100 % of humidity and blotted the solution for 5.5 to 6 s with blot force of either 19 or 20.

The data were acquired and analyzed as described before (Bromberg *et al*., 2020). Figures were prepared with PYMOL (The PyMOL Molecular Graphics System, Version 2.3.2, Schrödinger, LLC.), Excel (Microsoft Office 365 ProPlus, Microsoft), and Adobe Illustrator (Adobe Illustrator 2020, Version 24.0.1, Adobe Inc.). PYMOL includes an APBS electrostatic plugin (Baker *et al*., 2001) to which we provided an input generated with 6VSA.pdb and the server http://server.poissonboltzmann.org/ (Dolinsky *et al*., 2004). Electrostatic surfaces are displayed at +/10 k_B_T/e^−^ at full color saturation.

## 3. Results and discussion

Our data processing and analysis indicate that His-tag presence greatly changes preferred orientation patterns (Fig. 1a and 1b) of 5-fold symmetric coproheme decarboxylase particles. Without His-tag, these particles show very strong preferred orientation, with most (~40%) having their 5-fold axis oriented along the electron beam and perpendicular to the water-air interface. With His-tag, more particles rotate on their side, which significantly changes the patterns of preferred orientation; ~78% of the particles have their 5-fold axes oriented at ~116° or equivalently, at ~64° (if the opposite orientation of the symmetry axis is used) to the water-air interface (Fig. 1 and 2). The direction of preferred orientation can strongly influence 3D reconstruction; if it is aligned with the symmetry axis, it generates only one back projection orientation, while if it is at a high angle to the symmetry axis, it generates a symmetry number of back projections to the 3D reconstruction. Therefore, the change from one pattern of preferred orientation to another can have high impact, even if the orientations are not uniformly sampled.

**Figure 1.**
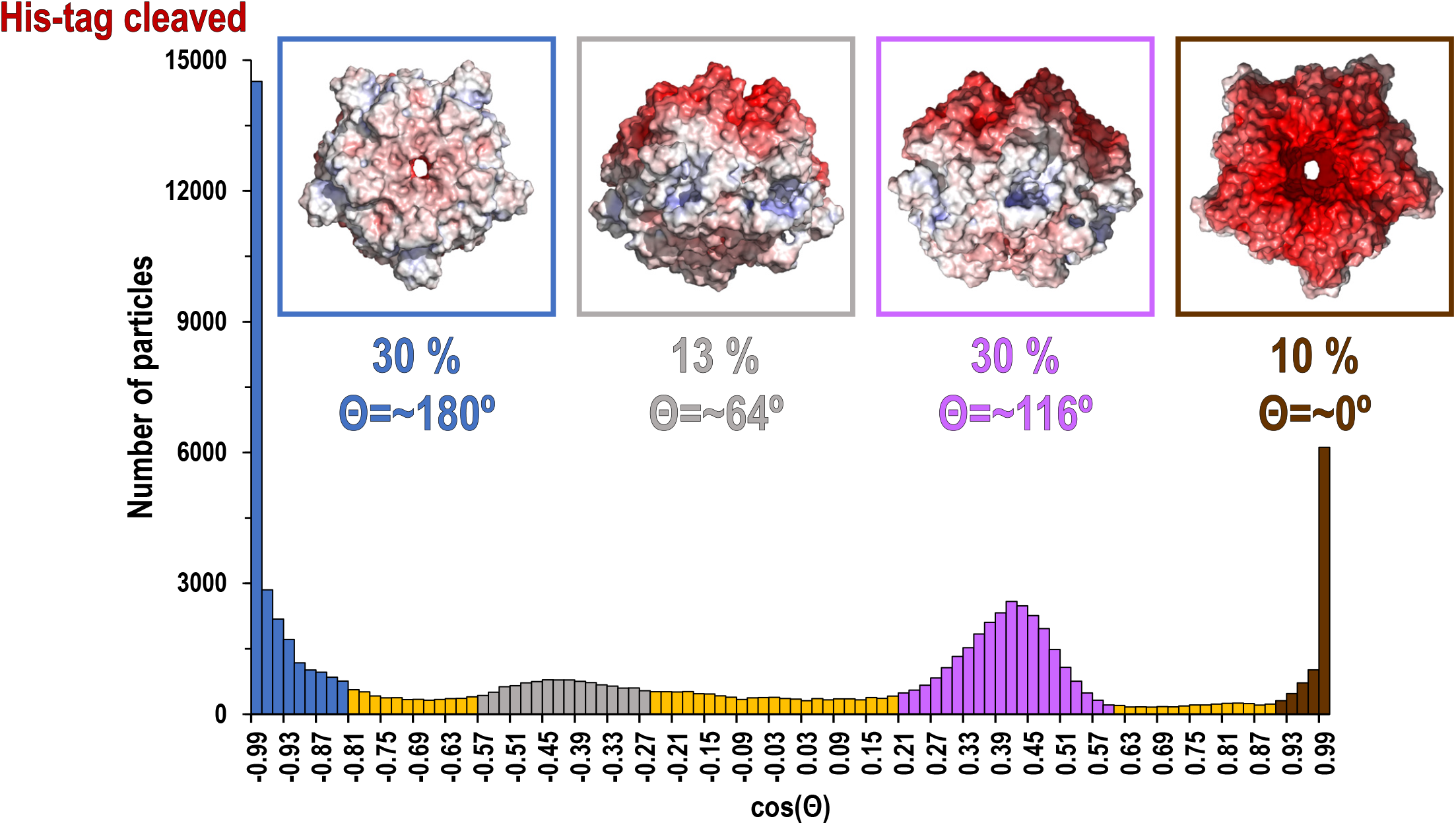
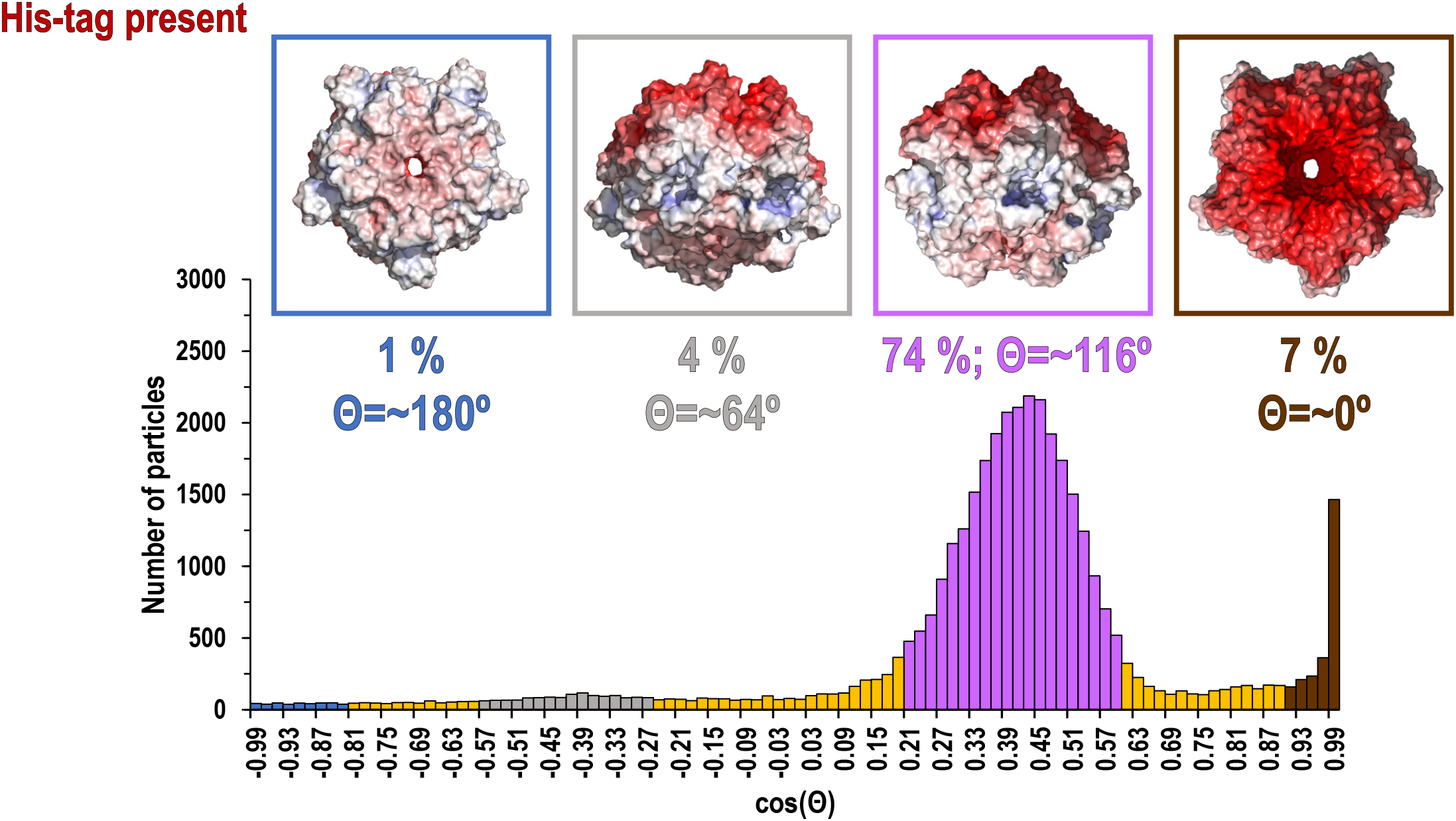
Histograms of distributions for particles of coproheme decarboxylase with His-tag cleaved (a) and His-tag uncleaved (b). The histograms show the number of particles as the function of cosine Θ, where Θ is the angle between the 5-fold axis of the particle and the direction of the beam. The map of the electrostatic potential for each orientation is shown above the specific peaks corresponding to the most frequent orientations.

**Figure 2.**
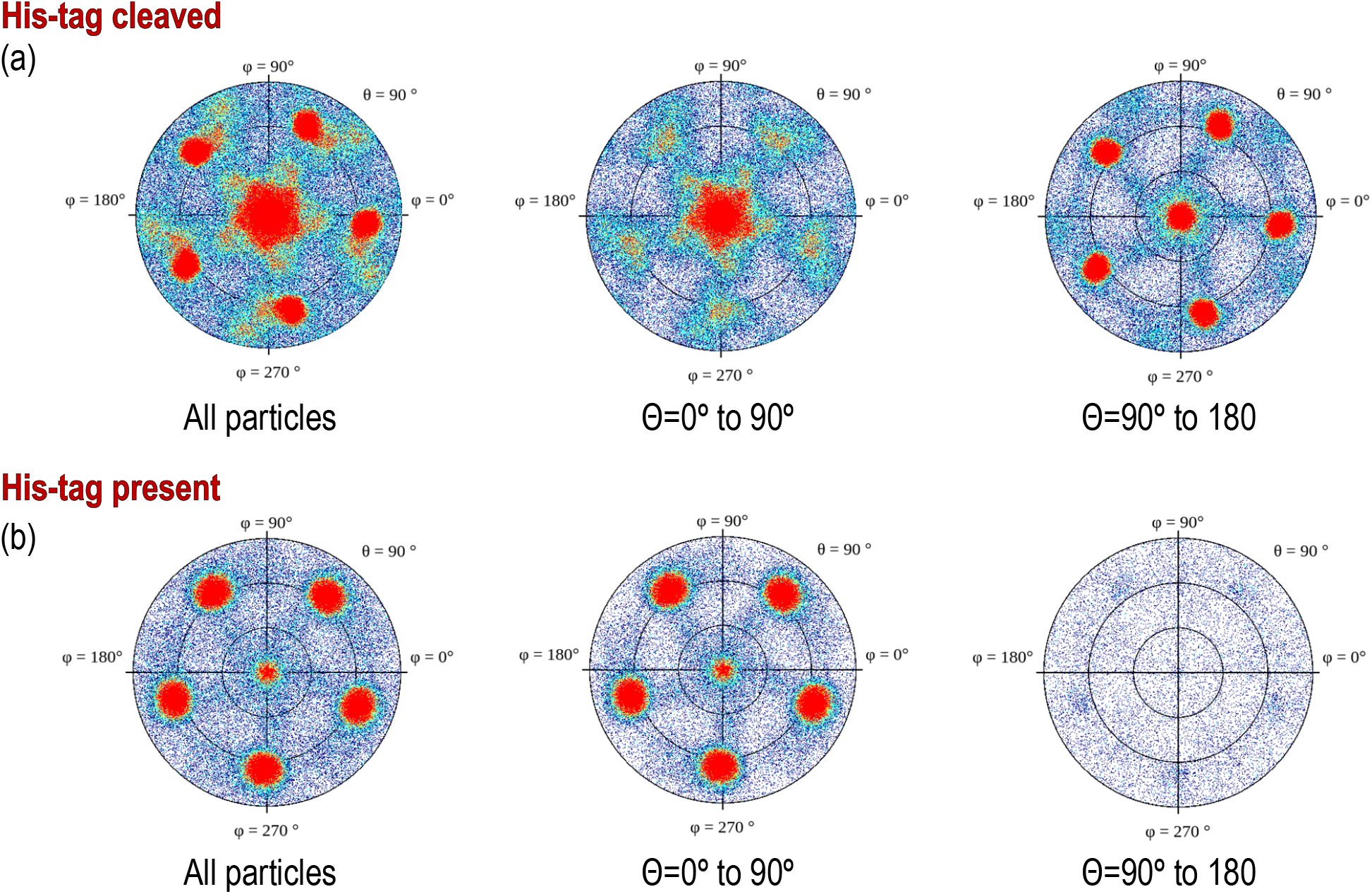
Preferred orientations (heat plots) for (a) His-tag cleaved protein and (b) His-tagged protein, shown in polar coordinates. To stress the difference in polarity of particles, we show on the left traditional plots for all particles, while the middle and right figures show (+) and (-) polarities (particles tilted away or toward the beam). The different polarities result from different angles of interactions with the water-air interface, but also may result from interactions with two separate water-air interfaces, which would symmetrize the histogram, as the cosine of the angle between the particle and the beam changes sign for the same geometric interaction with the water-air interface. We do not directly know how much these two effects contribute to our data.

2D orientation plots are not straightforward to interpret quantitatively. For this reason, we projected them on a single axis, which is the angle between electron beam and symmetry axis (Θ). Because our particles are polar, we can differentiate between parallel and anti-parallel orientations. The choice regarding which are considered parallel is arbitrary, but we keep it consistent in our figures. For such 1D orientation plots, uniform angular sampling would result in a sine modulation. We compensated for it by presenting the plots in uniform steps of cos(Θ). A random angular distribution will be flat in such a representation. The extremes in the range of cos(Θ) represent parallel and antiparallel to the beam orientation of particles. This representation is natural for particles with n-fold symmetry, but there is no natural choice of Θ for particles without symmetry.

The data show only the orientation with respect to the beam, including the polarity aspect. However, if we are interested in polarity of orientation with respect to the water-air interface, we need to consider the possibility of particles that bound to both water-air interfaces in a thin layer to be cryocooled. The strong polarity pattern, i.e. the observed asymmetry (Fig. 2), indicates that the frequency of binding to these two interfaces is very different. To precisely establish the apolar binding pattern for a single water-air interface would require tomographic reconstruction to isolate data arising from each interface. Past reconstruction tomographic data of particles trapped in a thin ice layer established that typically there is strong asymmetry in binding to top and bottom (relative to the beam direction) interfaces (Noble, Dandey, *et al*., 2018, Tan *et al*., 2014, Klebl *et al*., 2020), and this is fully consistent with our data. For polar particles, an advantage of our method is that it provides a quick and convenient characterization of asymmetry between two surfaces, and it also allows for retrospective analysis of cryoEM SPR data. The width of the peaks in our histogram (Fig. 1) result not only from the range of the angles between particles and the water-air interface, but also from variations in the tilt between the water-air interface and the beam. For typical grids, this is on the order of a few degrees (Noble, Dandey, *et al*., 2018) when data are acquired with at zero tilt angle. In addition to traditional 2D frequency plots, we also calculated 2D plots for two opposing polarities, to emphasize the significance of asymmetry of binding the water-air interface in two opposing directions (Fig. 2).

The significant change in patterns of preferred orientation between a protein with and without a His-tag prompted us to analyse 6VSA.pdb and 1T0T.pdb to identify possible reasons for the observed rearrangements.

Coproheme decarboxylase from *Geobacillus stearothermophilus* is a compact structure which has a strikingly asymmetric charge distribution, with only a small number of possible hydrophobic patches that would be attracted by the water-air interface (Fig. 3a). One of the patches is located at the bottom of the pentamer (Fig. 3a) and this patch most likely facilitates preferred orientation for proteins without His-tags (Fig. 1a). However, proteins retaining the His-tag had their N-termini extended by the sequence, **HHHHHHSSGVDLGTENLYFQSNA**, 23 amino acids in length, and originating from the MCSG7 vector. This part of the structure had no reconstructed density, as expected for parts of the chain that are disordered. However, this disordered sequence can extend up to ~3.25 Å per amino acid residue, so it can extend up to ~60-65 Å, which is more than the radius of the reconstructed part of the particle. The N-terminal tail can for example dynamically bind the flat, hydrophobic patch at the bottom of the pentameric assembly, and so preventing binding of this hydrophobic patch to the water-air interface. Alternatively, this tail may also bind directly to the water-air interface.

**Figure 3.**
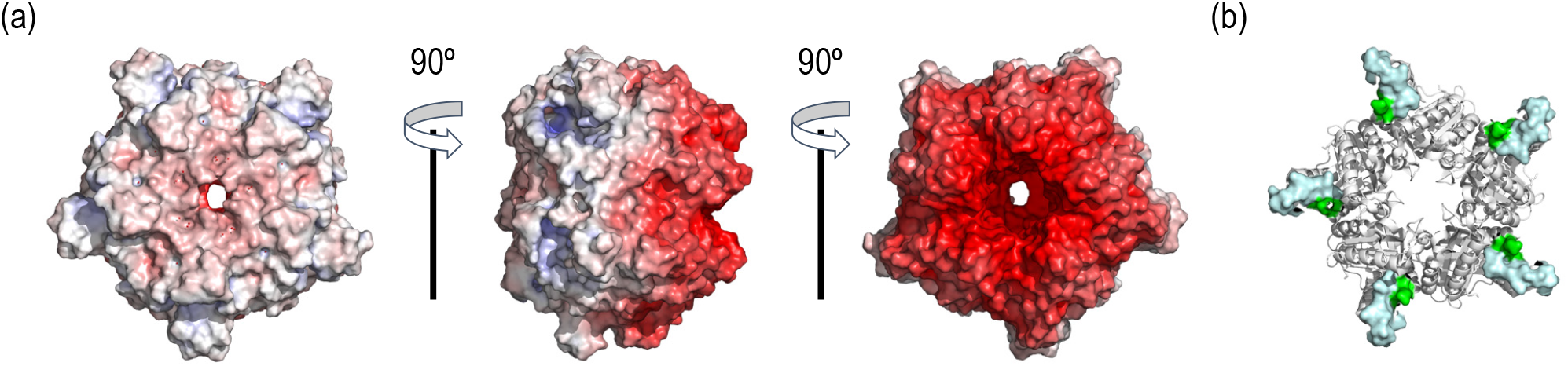
(a) The electrostatic potential map for 6VSA.pdb which represents the cryoEM reconstruction of the cleaved version of coproheme decarboxylase. (a) Three different orientations that stress a highly polarized charge distribution for this protein that explains observed patterns of preferred orientation. (b) The 1T0T.pdb X-ray structure of the same protein, in which the hydrophobic loop between 110-120 is ordered (light blue, surface representation). The loop is in proximity with the N-terminus (green, surface representation).

The second hydrophobic patch is located at a highly hydrophobic surface loop between amino acids 110 and 120 (**PAYSYVSVVEL**) (Fig. 3b). This region was ordered in the X-ray structure due to stabilizing interactions provided by the crystal lattice and a polyethylene glycol molecule, but in the cryoEM reconstruction (with and without His-tag) it is not visible. This unstructured region in our reconstruction is likely to serve as the water-air interface anchor and become partially unfolded (D’Imprima *et al*., 2019, Glaeser & Han, 2017), while the compactness of the rest of the structure prevents further unfolding. If the region were only unfolded upon binding to the water-air interface, then only 2 out of 5 loops in the pentamer would be affected, with the remaining 3 providing a weak but unambiguously visible contribution. The lack of this contribution indicates that the loop has multiple conformations, even outside of the water-air interface. This loop seems to direct the orientation of our particles, both for His-tagged particles and those without a His-tag. Additionally, the N-terminus of the ordered part, which is very close to the end of the His-tag, is also close to this surface loop. The much stronger presence of the side orientation may be due to the water-air interface interacting synergistically with both the hydrophobic part of the His-tag and the surface loop.

The two effects together: (1) the His-tag acting as a decoy obscuring the more hydrophobic, flat surface and (2) the linker between the N-terminus of the protein and the His-tag enhancing the hydrophobicity of the already hydrophobic loop create the observed distribution.

If retaining the His-tag allows for more general modulations of preferred orientation, then this suggests additional biochemical strategies for modulating interactions with the water-air interface and modulating the preferred orientation that is driven by these interactions: for example, modifying the length and the nature of the linker between the His-tag or other affinity tags, mixing tagged and untagged protein, attaching other decoy molecules, e.g. pegylation, or using reductive methylation of lysine residues (Means & Feeney, 1990, Rypniewski et al., 1993), to name just a few. Reductive methylation was used to change the pattern of hydrophobicity on the surface of proteins to promote their crystallization (Tan et al., 2014, Kim et al., 2008). One can expect that reductive methylation will have a similar impact on interactions with supports used in cryoEM SPR. Finally, one can also use a simple additive (e.g. 0.2 mM Ni_2_SO_4_) to adjust the state of His-tags in a labelled protein. This strategy was already successfully used in the case of membrane proteins, to change their associations with micelles (Rasmussen *et al*., 2019) and consequently change their aggregative properties.

Using tags as anchors and decoys to modify interactions with the water-air interface and consequently modulate patterns of preferred orientation offers an additional strategy to improve sample handling for cryoEM.

## Acknowledgements

We thank the Cryo-Electron Microscopy Facility (CEMF) at UT Southwestern Medical Center which is supported by grant RP170644 from the Cancer Prevention & Research Institute of Texas (CPRIT) for maintaining a Talos Arctica microscope.

## Author Contributions

RB, YG, DP, TE, MP, DB and ZO generated data; RB, DB and ZO analysed data; RB, YG, DP, TE, MP, DB, and ZO wrote manuscript.

## Conflict of interest statement

RB, YG, DB, and ZO are co-founders of Ligo Analytics. YG serves as the CEO of Ligo Analytics, RB and DP are currently employed by Ligo Analytics. ZO is a cofounder of HKL Research.

